# A bleb initiation model for chemotaxis suggests a role for myosin II clusters and cortex rupture

**DOI:** 10.1101/2020.03.16.993238

**Authors:** E.O Asante-Asamani, Daniel Grange, Devarshi Rawal, Zully Santiago, Derrick Brazill, John Loustau

## Abstract

Blebs, pressure driven protrusions of the plasma membrane, facilitate the movement of cells such as the soil amoeba *Dictyostelium discoideum* and other eukaryotes such as white blood cells and cancer cells. Blebs initiate or nucleate when proteins connecting the membrane to the cortex detach, either as a result of a rupture of the cortex or as a direct consequence of a build up in hydrostatic pressure. While linker detachment resulting from excess hydrostatic pressure is well understood, the mechanism by which cells rupture their cortex in locations of bleb formation is not so clear. Consequently, existing predictive models of bleb site selection do not account for it. To resolve this, we propose a model for bleb initiation which combines the geometric forces on the cell cortex/membrane complex with the underlying activity of actin binding proteins. In our model gaps, resulting from a rupture of the cortex, form at locations of high membrane energy where an accumulation of myosin II helps to weaken the cortex. We validate this model in part through a membrane energy functional which combines stresses on the cell boundary from membrane tension, curvature, membrane-cortex linker tension with hydrostatic pressure. Application of this functional to microscopy images of chemotaxing *Dictyostelium discoideum* cells identifies bleb nucleation sites at the highest energy locations 96.7% of the time. Sensitivity analysis of the model components points to membrane tension and hydrostatic pressure, all of which are regulated by myosin II, as critical to model predictability. Furthermore, microscopy reveals discrete clusters of myosin II along the leading edge of the cell, with blebs emerging from 80% of these sites. Together, our findings suggest a critical role for myosin II in bleb initiation through the formation of gaps and provides a predictive mathematical model for quantitative studies of blebbing.

**Author summary:** Eukaryotic cells such as white blood cells and cancer cells have been observed to move by making spherical herniation of their plasma membrane, referred to as blebs. The precise mechanism by which cells select locations around their boundary to initiate blebs is unclear. We hypothesize that blebs initiate at locations of high membrane energy where an accumulation of myosin II helps to rupture the cortex and/or detach linker proteins. We test this hypothesis by formulating a free energy functional representation of membrane energy to predict where blebs will initiate. The functional accounts for geometric forces due to membrane tension, curvature and membrane-cortex linker tension as well as hydrostatic pressure. Application of the functional to data from the soil amoeba, *Dictyostelium disodium*, identifies blebs at the highest energy locations over 90% of the time. Sensitivity analysis of model components points to membrane tension and hydrostatic pressure, all influenced by myosin II, as major forces driving bleb initiation. Additionally, we observe clusters of myosin II at locations of bleb initiation, further supporting its role in the process.

## Introduction

Chemotaxis, or chemically directed cell motility, is important for a variety of eukaryotic biological processes such as the development of an embryo, the search for pathogens by neutrophils, organ patterning, migration of fibroblasts to heal wounds and invasion of surrounding tissues by cancer cells [1–6]. An understanding of how cells chemotax will enhance our ability to intervene in these processes and improve the quality of life of our species. Traditionally, chemotaxis has been characterized by three actin-based motility structures filopodia, pseudopodia and lamellipodia [7]. These motility structures gained prominence in part because cells were predominantly observed while growing on a culture dish or crawling freely on the surface of a slide, all representative of a two-dimensional environment. Recently, observation of cells in their native three-dimensional environment, has uncovered a fourth motility structure, the bleb [8]. Blebs are pressure driven blister-like protrusions of the cell membrane. They are characterized by an initial detachment of the cell membrane from the cortex (nucleation), an expansion of the membrane into a spherical cap under force of flowing cytosol and a stabilization of the protrusion by reformation of the cortex [1,8–10].

Blebs have been studied from several viewpoints. They may be described as geometric objects (Euclidean) [11], the result of forces on a smooth manifold (differential geometry) [12–15], the result of physical force (pressure) [16–19], the result of cortical tension (related to pressure) [6] and the result of fluid dynamics [20, 21]. In spite of the substantial progress made in the study of bleb-based motility, the precise mechanism by which cytoskeletal proteins interact with biophysical and geometric forces to generate blebs and coordinate cell movement using these protrusions is still unclear. As a first step towards a comprehensive understanding of bleb-based motility we are interested in clarifying the interaction between proteins and forces leading to bleb nucleation or initiation. We take advantage of the genetic simplicity, ease of maintenance and chemotactic ability of *Dictyostelium discoideum,* a soil living amoeba, to study bleb-based motility [22].

Most researchers agree that blebs initiate when a small patch of membrane is detached from the cortex [1,8,15]. What differs among the theories of nucleation is the mechanism by which the plasma membrane detaches. One school of thought attributes the cortex-membrane detachment directly to hydrostatic pressure differential between the anterior and posterior ends of the cell [15,17,23,24]. The excess pressure detaches membrane to cortex binding proteins (linker proteins) and dislodges the membrane. In [15] a mathematical model is developed that uses sites of greatest tension in linker proteins to detect where the next bleb will form. This model is successful when *Dictyostelium discoideum* cells are subjected to high compression forces (2% agarose overlay) but does not perform as well when the compression force is lower (0.7% agarose overlay). This suggests that a different mechanism for nucleation may be at play when cells are subjected to low compression forces, necessitating a more universal marker of bleb nucleation.

Alternatively, initial membrane detachment is thought to be facilitated by local rupture of the actin cortex [25–27]. At these locations of cortex rupture, the membrane no longer has the support of the cortex. The force of cytoplasmic flow is then capable of breaking the adhesive bonds and detaching the membrane. Recent work has attributed the degradation of the cortex to the action of myosin II [2,16], yet neither a direct evidence of cortex rupture in live blebbing cells nor of the direct involvement of myosin II is provided. Furthermore, it is reasonable to assume that cells will use cortex rupture to assist nucleation under low compression force, where intracellular pressure may be insufficient. Since linker detachment via cortex rupture may not require linker proteins to be maximally stretched, the use of locations of maximal linker stretching as a locator of nucleation sites may be insufficient. This may very well explain the poor performance of the predictive model in [15] when cell are under low compression force.

In this study, we introduce a model of bleb nucleation that unifies the two existing points of view and provides a mechansim for cortex rupture. Any shape has an energy cost and when unconstrained will seek to transform into a configuration with lower energy [28,29]. We expand this idea and propose that blebs occur at locations of high boundary stress, where activities leading to the rupture of the cortex or stretching of linker proteins increases the local energy. Once the linker proteins which hold the membrane to the cortex are detached, the free membrane expands into a mostly circular configuration with nearly constant energy. Hence, there are no longer high energy spikes and the cost of maintaining the shape is lowered. A mathematical model which represents the membrane energy as a Helfrich free energy functional [30,31] is developed and used to test this hypothesis. The model extends the energy functional described in [15] by including a spatially varying hydrostatic pressure term. This term accounts for energy contribution from local myosin II activity in the cortex, which has been suggested to contribute to cortex rupture [2,18]. To validate our model predictions, we present a geometric marker for detecting nucleation sites directly from microscopy images. In order to speed up the data collection process, we introduce an algorithm for automatic detection of blebs from processed microscopy images. Our mathematical model when applied to data from *Dictyostelium discoideum* cells successfully predicts over 90% of observed nucleation sites, about 70% of which show gaps in the cortex. To clarify the role of myosin II in bleb nucleation, we perform localization experiments which suggest a critical role for the motor protein in the formation of cortex gaps.

## Materials and methods

### Strain and culture conditions

All *D. discoideum* cells were grown axenically in shaking culture in HL5 nutrient medium with glucose (ForMedium) supplemented with 100 *μ*g/mL penicillin and streptomycin (Amresco) at 150 rpm at 22°C. LifeAct-GFP expressing Ax2 wild type cells were grown in 4-20 *μ*g/mL G418 (Geneticin). For LifeAct-RFP expressing cells, 50 *μ*g/mL Hygromycin B was used and 4 *μ*g/mL G418 for myosin-GFP expressing cells. Cells were starved for cAMP competency on filter pads using the method described in [32].

### cyclic-AMP(cAMP) under agarose assay

Two cAMP under agarose assays were used as described in [32], with minor modifications, to mimic the confinement conditions cells experience in their natural environment. The first under agarose assay was used to collect data on blebbing cells for validating the nucleation site predictions of the membrane energy functional. For this assay, wild type cells crawled under a 0.7% Omnipur agarose (EMD Millipore) gel that was laced with 1 mg/mL of 70,000 MW Rhodamine B isothiocyanate-Dextran (Sigma-Aldrich). In this way, the membrane could be observed as a shadow within a red background created by the Rhodamine-Dextran (see [32]). In the second assay, used for myosin II localization, the agaros gel was not laced with Rhodamine B isothiocyanate-Dextran. Rather, the membrane position was identified using a bright field light source. This was done to permit the identification of the cortex through the expression of LifeAct-RFP.

### Image acquisition

Imaging data for the under agarose assay 1 were collected using a Leica DMI-4000B inverted microscope (Leica Microsystems Inc.) mounted on a TMC isolation platform (Technical Manufacturing Corporation) with a Yokogawa CSU 10 spinning disc head and Hamamatsu C9100-13 EMCCD camera (Perkin Elmer) with diode lasers of 491 nm, 561 nm and 638 nm (Spectra Services Inc.). Images for the under agarose assay 2 were collected using a Perkin Elmer UltraView ERS spinning disk confocal microscope equipped with the same lazers used in the first assay. For the first assay, LifeAct-GFP and RITC-Dextran were excited using the 491 nm and 561 nm lasers. Cells were imaged for 30 seconds using either 80x magnification (40x/1.25-0.75 oil objective with a 2x C-mount) or 100x 1.44 oil immersion objective. Data collected using both GFP and RITC channels resulted in a frame rate of 1.66 seconds whereas data collected using only GFP resulted in one frame rate of 0.800 seconds. For the second assay the 491 nm and 561 nm lasers where used to excite myosin-GFP and LifeAct-RFP. A bright field light source was used to illuminate the membrane. *Image J* was used to adjust brightness and contrast of the images, which were then imported into our Mathematica based geometric system for edge detection, boundary digitiation and automatic bleb detection.

### Edge detection and boundary digitization

We modified an in-house geometric platform [32] which extracts the boundary of photographic images and renders them as objects in Cartesian space. Our modified algorithm, in addition to the procedure outlined in [32], thins the cell boundary to a width of one pixel (uses the *Thinning* function in *Mathematica*), implements a more efficient algorithm for ordering boundary points, orients the boundary, and finally applies a Gaussian convolution to smooth the boundary. The digitization of the boundary into equally spaced points using B-splines, is also implemented more efficiently. These modifications were necessary to speed up the data processing and reduce jagged boundaries, which can throw off our energy calculations.

### Ordering boundary nodes

In our previous work [32] we applied the *FindShortestTour* function in *Mathematica* to order the coordinates of boundary points. For cells with complex boundaries, this function often failed to yield a reasonable ordering of the points. Additional attempts to remedy this by ordering smaller segments of the boundary was very time consuming. For our current application, we have completely eliminated the use of the *FindShortestTour* function and have instead implemented a straightforward algorithm to order boundary points. Let 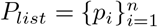 be the set of boundary points, beginning from the first point in the list, *p*_1_ = (*p*_1*x*_,*p*_1*y*_) we obtain a set of candidates for the next ordered point, 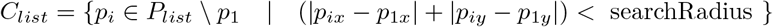. The next ordered point after *p*_1_ is then chosen to be the point in *C_list_* closest *p*_1_ in Euclidean distance. *P_list_* is reset to *C_list_* and the process is repeated (see Fig. 1A). The use of the Manhattan distance instead of the Euclidean distance to obtain *C_list_* allows us to quickly search through the boundary points. Our revised algorithm always yields a proper ordering of boundary points and took on average 0.36 seconds per frame.

**Fig 1.**
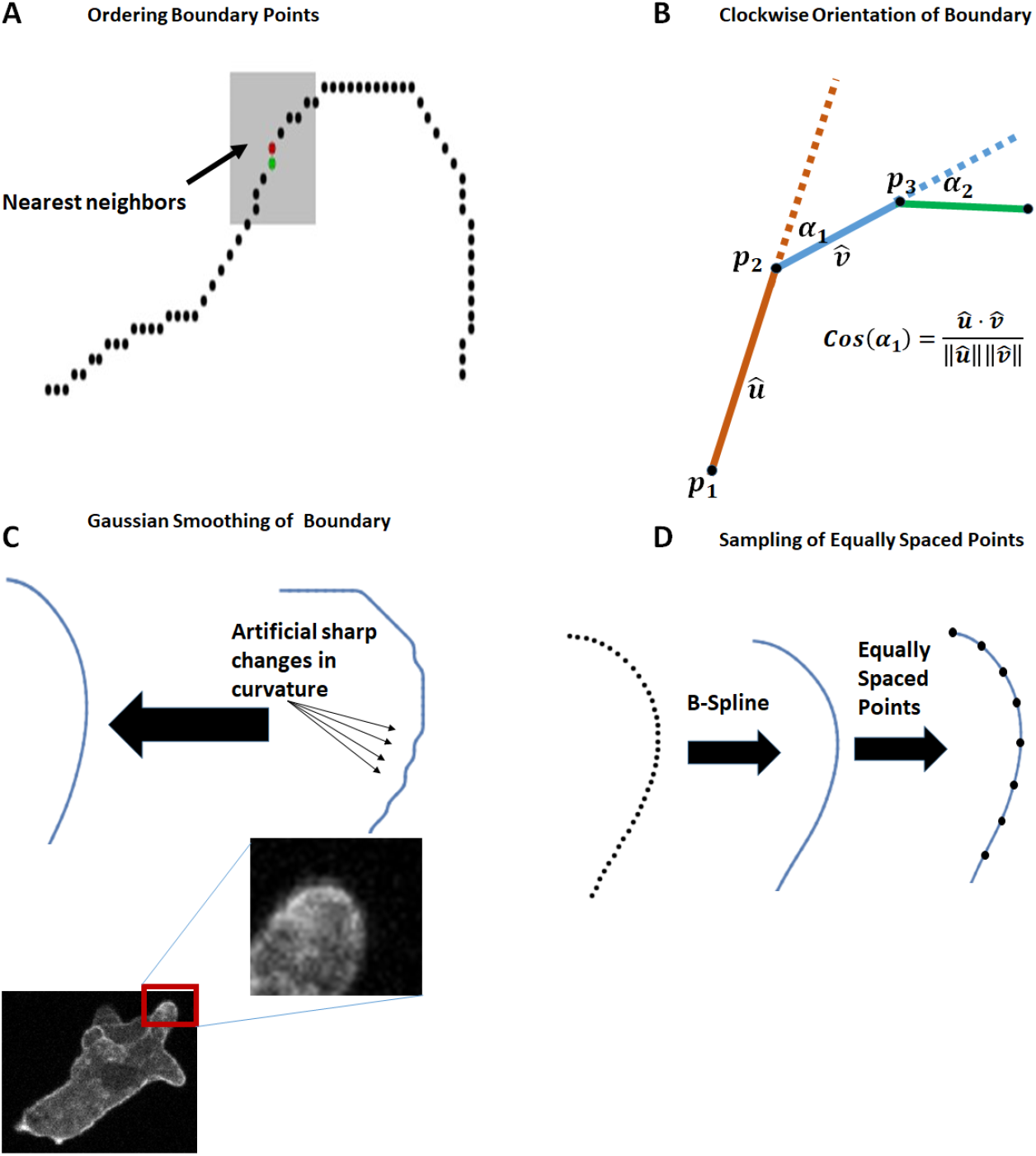
Cell boundary digitization, orientation and smoothing. The techniques used to digitize, orient and smooth the boundary of cells are shown. A) A scatter plot of boundary points is shown with a shaded region denoting the nearest neighborhood of the current boundary point (shown in red) obtained using the Manhattan distance metric. The next ordered point (shown in green) is subsequently obtained as the closest point (in Euclidean distance) to the current point. B) The turning angles *α*_1_ and *α*_2_ for the triplicate *p*_1_,*p*_2_,*p*_3_ are shown. C) The edge detected from a portion of the cell shows artificial sharp changes in curvature which are resolved using Gaussian smoothing. D) Equally spaced points are generated from a B-spline representation of the ordered, oriented boundary point.

### Clockwise orientation of boundary nodes

Our purpose in ordering the boundary nodes is to make it easier to automatically detect blebs. We chose to fix a clockwise orientation for all boundary points, *P_list_* as follows. Let *P_convex_* denote the boundary of the convex hull of *P_list_*, a portion of which is shown in Fig. 1B. For each triplicate of points, {*p*_1_,*p*_*i*+1_,*p*_*i*+2_} we set the vectors *û* = *p*_*i*+1_ – *p_i_* and 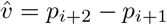 and calculate the turning angle, *α* using the formula

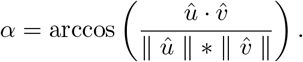

The ordering of the triplicate is clockwise if 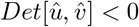 and counterclockwise otherwise. The turning angle is assigned a positive or negative sign depending on the orientation of the triplicate. We consider the ordering of *P_list_* to be clockwise if sum of all turning angles is –2*π* and counterclockwise otherwise. In the event of the latter, we reverse the order of *P_list_* to achieve a clockwise orientation.

### Gaussian smoothing of boundary nodes

Our calculation of membrane energy is dependent of an accurate representation of boundary curvature. Quite often, the cell boundary obtained from the edge detect process had some jaggedness not present in the original image (see Fig. 1C). To recover biologically relevant boundary curvature, we perform a discrete Gaussian convolution on boundary points by setting each point, 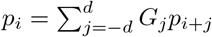 with weights *G_j_* sampled from a standard normal density function with standard deviation, *d*. In Fig. 1C, we show how the smoothing process removes artificial curvature artifacts in the cell boundary.

### Generation of equally spaced boundary nodes

Our membrane energy functional assumes a smooth parameteric representation of the cell boundary from which derivatives are calculated. We achieved this paramterization using B-splines, as described in our previous work [32] and employed finite differencing to approximate the derivatives. In order to simplify the implementation of finite difference schemes and reduce the local truncation error [33], we sought a partitioning of the boundary into equally spaced points. In this work, we optimized the boundary discretization procedure described in [32] to reduce the computer processing time.

A B-spline representation of the cell boundary, using *n* oriented boundary points, 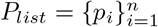 requires fitting *n* – 3 segments of twice continuously differentiable curves with the *i^th^* segment, *σ_i_*(*t*): [0,1] → ℝ^2^ constructed to lie in the convex hull of four consecutive (guide) points {*p_i_,p*_*i*+1_,*p*_*i*+2_,*p*_*i*+3_}. The result is a cubic polynomial formed from the convex combination of the guide points,

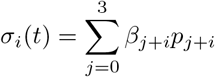

using basis functions

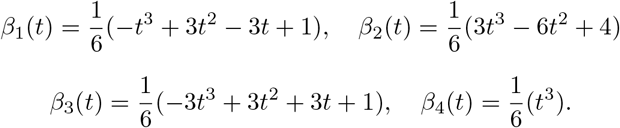

Let 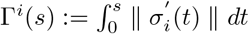 then the total distance along the cell boundary can be expressed as

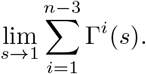

In our previous work, we partitioned the boundary into segments of length 0.5 pixels using the following iterative process:

1. While |Γ(*s*) – 0.5| > *tol*
2. Set *s* = *k*Δ*s*
3. Numerically integrate Γ(*s*) using the *NIntegrate* function in Mathematica.
4. Set *k* = *k* + 1 and go to step 1.

With additional lines of code to translate residual segment distances to the next segment the process took on average 41.23 seconds per frame and 13 minutes for each cell(20 frames). Much of the computational time was spent adjusting the limits of integration until an appropriate limit was obtained to generate the desired segment length. We circumvented this costly search process by directly integrating a Lagrange polynomial interpolation of the integrand, 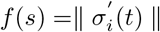 once and solving for the desired upper limit using a root search algorithm such as Secant method. For 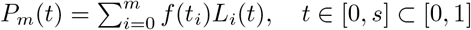 an *m^th^* degree Lagrange polynomial with Lagrange basis functions 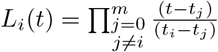, we set 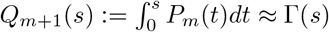, and determined a segment with length 0.5 pixels by solving the nonlinear equation *Q*_*m*+1_ – 0.5 = 0. The revised algorithm, took 0.57 seconds per frame which is a substantial improvement over our previous effort. The process is illustrated in Fig. 1D.

### Automatic detection of blebs

The detection of blebs and subsequent analysis of their characteristics such as energy, nucleation point, area and boundary curvature can be a very time consuming process. This is in part because most statistical tests require large sample sizes to yield relevant outcomes. In the case of calculating bleb area for example, one has to visually distinguish blebs from other actin-based protrusions and then isolate the boundary of each bleb for use in software packages such as *Image J* or *Mathematica.* In our application, we image cells for about 30 secs at an average frame rate of 1.66 secs. This generates between 18 and 20 frames per cell. To draw reasonable statistical conclusions, using between 20 and 40 cells, we would have to analyze about 160 blebs, visually detected from up to 800 frames.

To reduce the time and energy required to detect blebs and improve the accuracy of measurements made on those blebs, we have developed an algorithm that automatically identifies blebs from a time-lapse sequence of discretized cell boundary profiles *(Equlist*) and calculates desired bleb characteristics. Once the discrete, equally spaced, oriented boundary profiles are inputed, the algorithm first identifies all protrusions using consecutive *Equlists*. The remaining parts of the algorithm utilize geometric properties of blebs to distinguish them from actin-based protrusion such as pseudopods (see Fig. 2). This is the first publicly available algorithm for detecting and analyzing blebs. We expect this computational tool to greatly enhance research into bleb-based chemotaxis. In the subsections that follow we present the mathematical constructs used to accomplish this automation.

**Fig 2.**
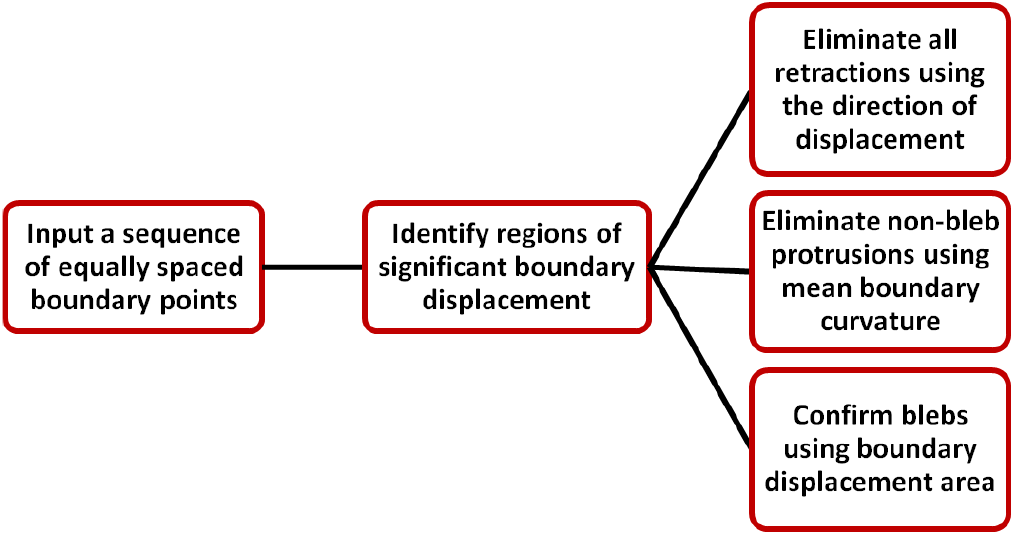
A flowchart showing the methodology for detecting blebs from a time-lapse of discretized cell boundaries.

### Detecting boundary protrusions

The set of distances between consecutive cell boundaries, taken along the normal direction, define a *displacement profile* (Fig. 3A). We refer to the boundary points corresponding to the first cell image as the *prior cell* and use *post cell* for the second cell image. For each boundary point on the prior cell, the displacement is given a positive or negative sign depending on whether the point of intersection with the post cell lies in the exterior or interior of the prior cell. A *protrusion* is defined by a set of points whose displacement exceeds an upper threshold, *PU_thresh_*.

**Fig 3.**
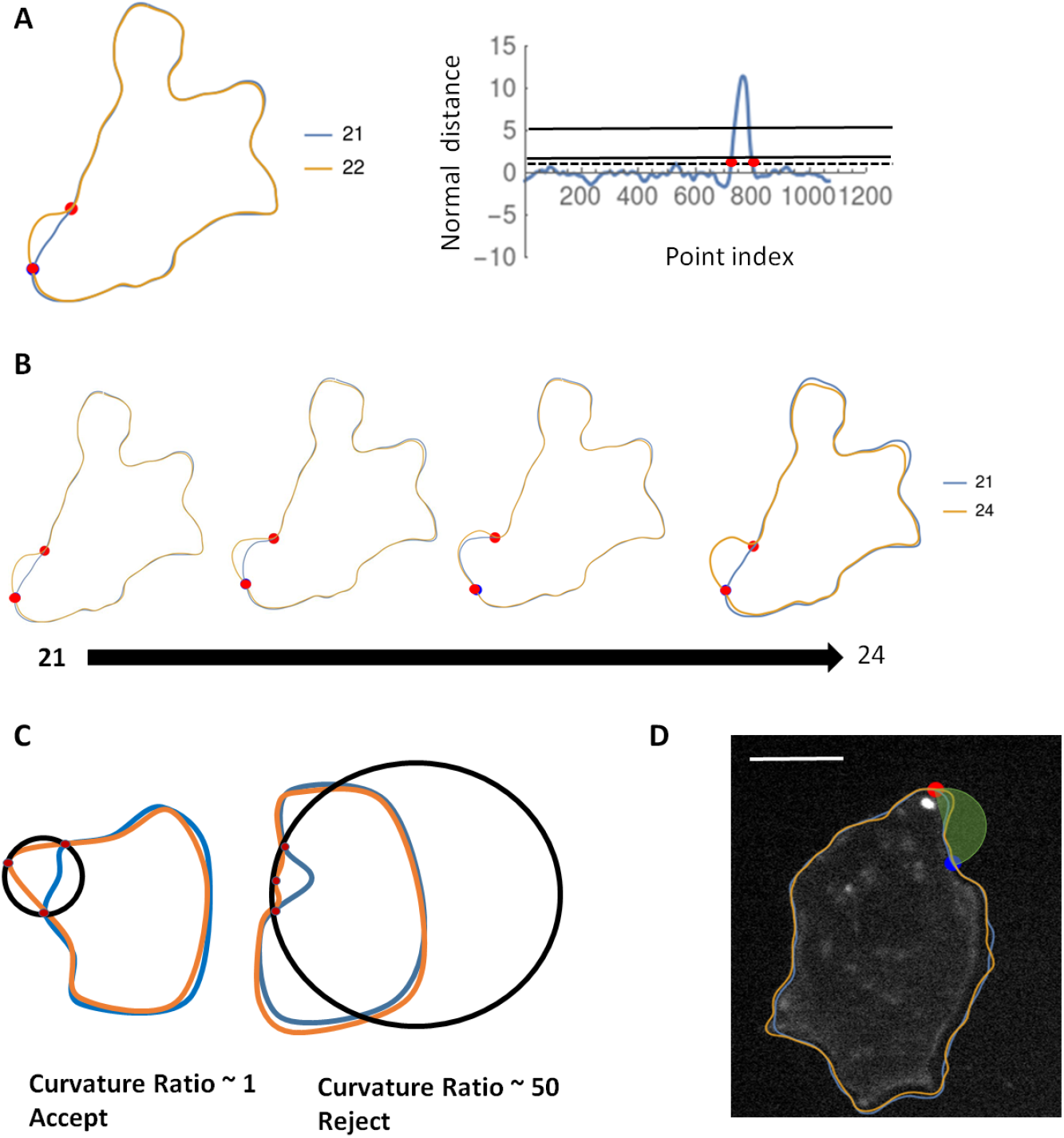
Computational scheme used to detect blebs from cell boundary profiles. A) Two successive boundaries of a chemotaxing D. discoideum cell are shown (left image) with the prior cell in blue and post cell in orange. The normal displacement between consecutive cell boundaries is used to generate a protrusion profile(right image) from which the bleb shoulder points (red dots) are identified. The two horizontal lines denote the *PU_thresh_* and *PL_thresh_*. B) Successive frames over which a bleb expands are combined to obtain the complete bleb shape. C) The mean curvature of the bleb boundary relative to a fitted circle is used to distinguish blebs from pseudopods. D) The area enclosed by the bleb boundary relative to the area of a fitted semi circle is used as a final confirmation of blebs. Scale bar is 5*μ*m

### Detecting shoulder points

For blebs, we refer to the points on the cell boundary where membrane peeling halts as *shoulder points*. These points should ideally be present in the *Equlist* corresponding to both the prior cell and post cell. Since the bleb boundary is usually split between two successive frames, the actin scar on the prior cell and the detached membrane on the post cell, the shoulder points serve as reference points for accurately defining the boundary. To determine these shoulder points, we first identify points in the displacement profile above *PU_thresh_* for which forward movement results in a point below *PU_thresh_*. Searching forward and backward, the shoulder points are identified as the first set of points that lie below a designated *PL_thresh_* in either direction (See red markers in Fig. 3A).

### Identifying the resting shape of protrusions

Since blebs develop over multiple frames, it is necessary to identify protrusions not only between successive frames, but by connecting frames over the full duration of bleb expansion. In order to identify the terminal frame of a particular bleb, we keep track of the extent to which the initial shoulder points are pushed outwards along the cell boundary. In particular, assume a clockwise ordering of boundary points and let *u*_1_, *u*_2_ be the indices of the left and right shoulder points for a protrusion on a prior cell and *v*_1_, *v*_2_ be the corresponding shoulder point indices on a post cell. Then the shoulder points correspond to the same protrusion if *v*_1_ < *ϵ* * (*u*_2_ – *u*_1_) and *v*_2_ > *ϵ* * (*u*_2_ – *u*_1_), where *ϵ* < 1 is determined experimentally. The bleb boundary is then defined as a list of points between *v*_1_, *v*_2_ on the initial frame (actin scar) joined with the points between *v*_1_, *v*_2_ on the terminal frame. See Fig. 3B for an illustration.

### Isolating blebs from other actin-based protrusions

Up until this point, the discussion has been focused on identifying protrusions, with no ability to distinguish between blebs and pseudopods. To make this distinction we use a two fold test which takes advantage of the geometric characteristics of blebs. First, since blebs are mostly spherical, we compare the average curvature of the leading edge of the protrusion (boundary minus actin scar) to the curvature of a circle fitted to the shoulder points and the midpoint of the boundary. We expect the curvature ratio to be close to one for blebs and further from one for pseudopods and other non-bleb protrusions. In Fig. 3C the curvature ratio is used to accept a protrusion as a bleb. Secondly, we compare the protrusion area to half the area of a fitted circle (Fig. 3D). Once the protrusions are determined to be blebs from these tests, the program gives the user an opportunity to make the final call by presenting microscopy images (cortex and membrane) corresponding to the initial and terminal frames of each identified bleb.

### An energy functional for predicting bleb nucleation sites

When under compression forces, such as experienced during multi cellular development, *D. discoideum* cells flatten and can be considered two dimensional. Thus our cell boundaries are represented as one dimensional smooth curves. We represent the membrane energy as a Helfrich bending energy functional [30] with additional energy contributions from tension in the linker proteins and local hydrostatic pressure,

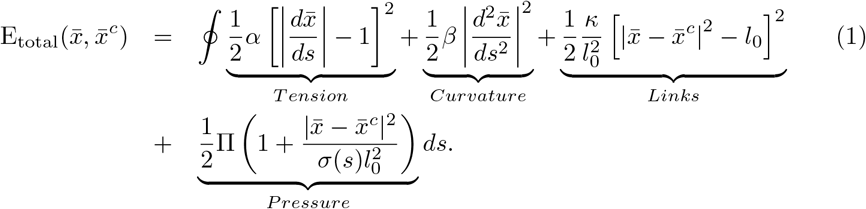

The membrane and cortex configuration, parameterized with respect to arclength *s*, are denoted by 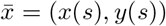 and 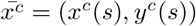 respectively. The first two terms refer to the energy contribution from membrane stretching and bending with membrane stiffness, *α*(*p_N_*/*μm*) and bending rigidity, *β*(*p_N_*/*μ_m_*). The third term is energy associated with tension in the linker proteins by modeling them as linear elastic springs with spring constant *κ*(*p_N_*/*μm*). The resting linker length, for which there is no tension, is denoted as *l*_0_ (*μm*). These three energy components are similar to the model in [15]. We introduce, in our fourth term, a spatially varying hydrostatic pressure term with ambient pressure Π(*P_a_*) and model dependent parameter *σ*(*s*). The values of *σ*(*s*) are computed so that the ratio 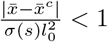. Our motivation for this term, is to develop a more biologically relevant model.

The value of our pressure term reflects the local activity of myosin II in the cell cortex. Local pressure increases as a result of cortex contraction by myosin II [2,34,35]. In constrast to [15], we take the point of view that equilibration of this pressure is not perfectly efficient, resulting in short lived spatial variations on a time scale consistent with bleb nucleation. This is supported by experimental work in [36], necessitating recent poroelastic formulations of the cytoplasm [20, 21]. Since cortex contraction by myosin II will stretch linker proteins, we use changes in the linker protein lengths as a measure of the level of local hydrostatic pressure. Our model adjusts the ambient global pressure Π by an amount proportional to the length of the linker protein, imposing a higher pressure in locations with more stretched linker proteins.

Whereas similar models of hydrostatic pressure have been used to describe bleb expansion [14, 37], we are the first group to use it in a quantitative study of bleb nucleation. In accordance with our hypothesis that blebs initiate at locations of high membrane energy, we restrict the cell boundary to the region between the endpoints of the newly formed bleb (bleb neck), and select the highest energy point as the most likely bleb nucleation site.

Blebs nucleate when proteins connecting the membrane to the cortex detach, either as a result of a rupture of the cortex or as a direct consequence of a build up in hydrostatic pressure [1]. In the model presented in [15], locations of maximal linker extension was used as a predictive marker for nucleation. Since a rupture of the cortex does not necessitate streching of the linker proteins, this marker may not adequately predict nucleation via cortex rupture. An advantage of using high energy to detect nucleation sites is that we are able to capture both an increase in tension due to stretching of the linker proteins as well as build of cortical tension preceding cortex rupture. Thus providing a more general predictive marker for nucleation.

We propose that blebs initiate at points of high boundary stress. This hypothesis, unlike previous models, permits an unbiased detection of bleb nucleation whether through linker detachment or gap formation. The energy functional captures the stress on the cell membrane which leads us to consider points with maximal energy value, a quantitative measurement, as the preferred means of initiation detection. In order to use the energy functional we needed to identify the location of the membrane and cortex across multiple frames, discretize their respective configurations and finally feed that information into a discretized version of the functional.

First, we determined the cortex (labeled with LifeAct-GFP) location using a Cany edge detect utility in Mathematica, fit a cubic B-spline to the x-y coordinates and then generated equally spaced points along the B-spline. This produced an approximate arclength parameterization of the cell cortex, 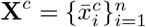 We then initialized the membrane to be an equidistant, outward normal, projection of the cortex, **X**^0^. The separation distance, 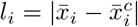 was set to the experimentally determined resting length of the linker proteins (0.04 *μ*m [12]). Following from the principle of least action, we determined the true membrane location to be the configuration 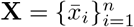 which minimized the discrete energy functional, 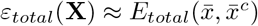, given by

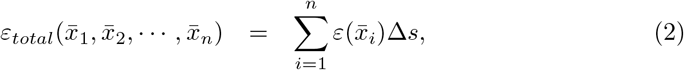

with the pointwise energy, 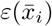, defined as

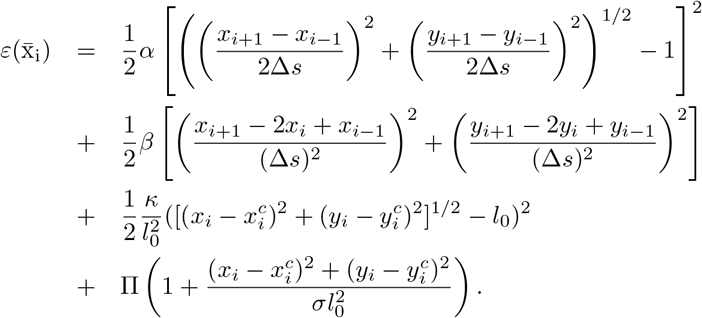

Here, the first and second derivatives in the continuous energy functional have been discretized using the first and second central finite difference schemes respectively. Δs is the designated fixed arclength interval.

Once the membrane location had been determined, we evaluated the pointwise energy within the bleb shoulder points and identified the location with maximum energy as the most likely nucleation site.

### A geometric marker for identifying bleb nucleation sites

In order to evaluate the predictive power of our energy model, we needed to be able to identify bleb initiation sites on chemotaxing cells, directly from microscopy images. For clarity, these initiation sites correspond to locations on the actin scar (old cortex between bleb shoulder points) where the membrane first detached from the cortex. To accomplish this, we have developed a geometric marker, that uses the final bleb shape to predict the initiation site. These sites are then compared with the predictions from the energy functional.

Since bleb expansion is driven primarily by fluid flow, it is reasonable to assume that the location on the newly formed cortex/membrane complex, furthest from the actin scar, will be contained in the membrane patch that initially detached from the cortex at initiation. We determine this location, the furthest extent of the bleb, on the newly formed cortex as the maximum perpendicular distance from the cortex to a line segment connecting the bleb shoulder points (bleb neck) (Fig. 4B). Once identified, the furthest extent is mapped to the actin scar, by extending its projection to the bleb neck. The point of intersection between the extension and the actin scar is chosen as the nucleation point (Fig. 4C). In Fig. 4(D,E) we show an application of the furthest extent to identify nucleation points in two chemotaxing *D. discoideum* cells.

**Fig 4.**
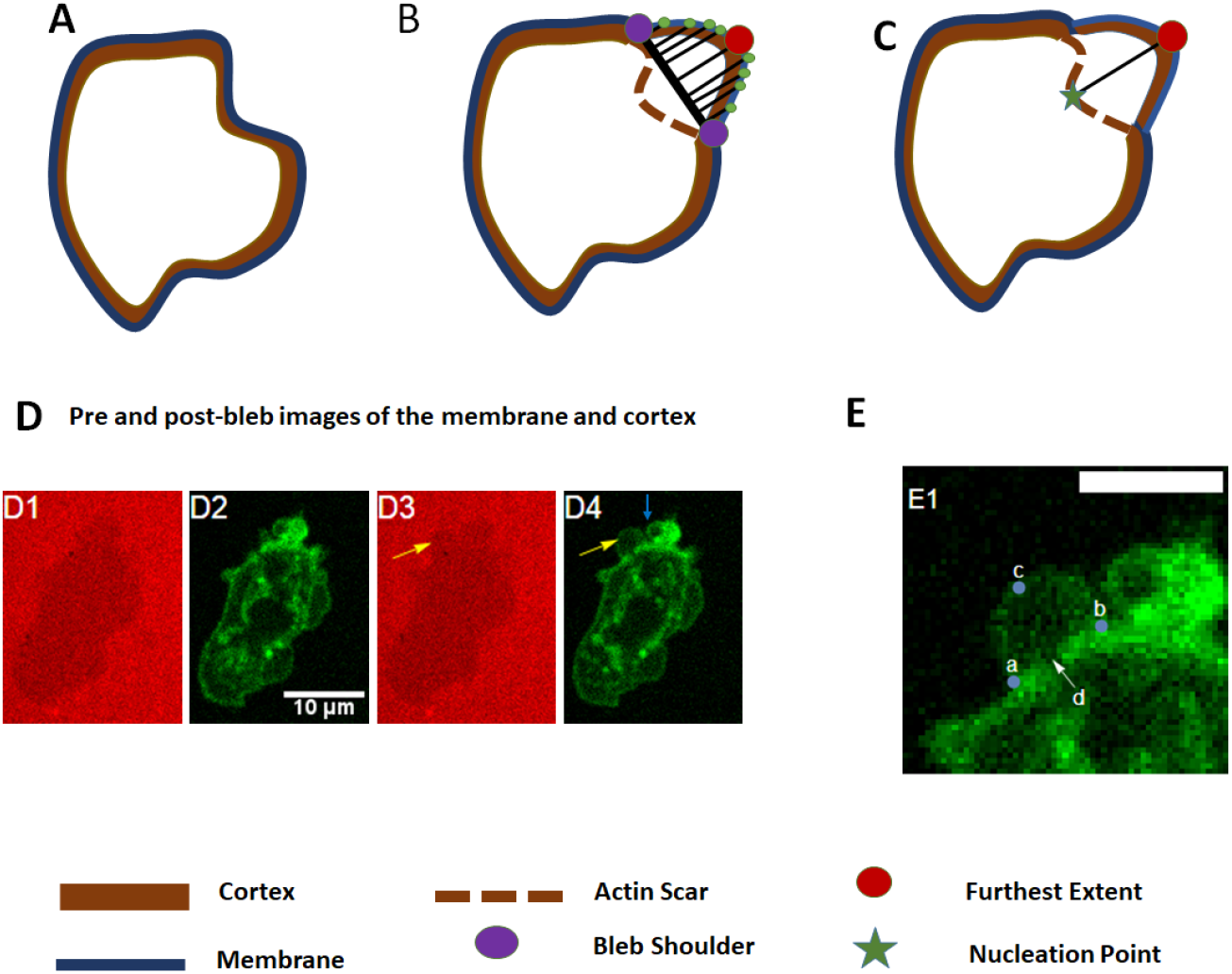
The furthest extent of the bleb identifies nucleation sites. A set of cartoons showing the methodology for determining the furthest extent of a bleb with application of the marker to microscopy images. A) A cell before bleb initiation (pre-bleb image). B) After the bleb initiates and stabilizes (post-bleb image), the perpendicular distance between points on the fully formed bleb and a line segment connecting the bleb shoulder points (bleb neck) is used to determine the furthest extent. C) The furthest extent is projected onto the actin scar to identify the nucleation point. D-E) the furthest extent is applied to identify nucleation sites in chemotaxing *D. discoideum* cells expressing lifeAct GFP. The B-spline representation of the pre-bleb cell boundary (blue curve) and post-bleb cell boundary (orange curve) are overlayed with the images. Scale bar is 10 microns. D) Pre-Bleb images of the cell are shown in D1 and D2. D1 is the membrane image and D2 is the corresponding cortex image. The post-bleb images are D3, membrane and D4, cortex. The arrows on D3 and D4 identify the newly formed bleb. Note there is no bleb in D1 and D2. E) Finding bleb features in order to infer nucleation in the pre-bleb image. Scale bar is 5 um. Images E1 shows the cortex just after formation of the bleb. We identify the bleb shoulder points a and b, the furthest extent of the bleb is denoted by c. A cortex gap is identified by d.

## Results

### The membrane energy functional robustly predicts bleb nucleation sites

Performing this experiment with model parameters *α* =17 *pN/μm, β* = 0.14 *pN/μm, κ* =10 *pN/μm*, Π = 81 *Pa* for 109 blebs, we were able to correctly predict nucleation sites in 105 cases (96.8%). These data suggest that the energy function has a high predictive ability for identifying bleb nucleation sites, free from the limitations of the human based process.

**Table 1.** Results from nucleation point detection using ROI framework.

| Line |  | Count | Percentage |
| --- | --- | --- | --- |
| 1 | Total number of cells | 33 |  |
| 2 | Total number of blebs | 142 |  |
| 3 | Confirmed nucleation points from ROI | 109 | 76.8% |
| 4 | Confirmed gaps | 79 | 72.5% |

It is reasonable to expect a degree of measurement error in the values of the model parameters, *α, β,κ*, Π. Ideally, such error should not significantly change our model predictions, if they are to be considered reliable. We investigated the robustness of our energy model using five tests. In the first test, we allowed each parameter to vary by 20% of their experimental value, generated 1000 uniform random samples from this interval and used each parameter set to predict a chosen nucleation site. The Euclidean distance between the predicted site and the validated site (using ROI framework) was used to measure the level of variation in model output. The subsequent tests were performed by varying model parameters by 40%, 60%, 80% and 100% of their measured valued. From Fig 5A, there is effectively no change in model predictions for up to 60% variation in their measured values. Since experimental error is generally not expected to exceed 40% of measured value, we conclude that the model predictions are robust to measurement error.

**Fig 5.**
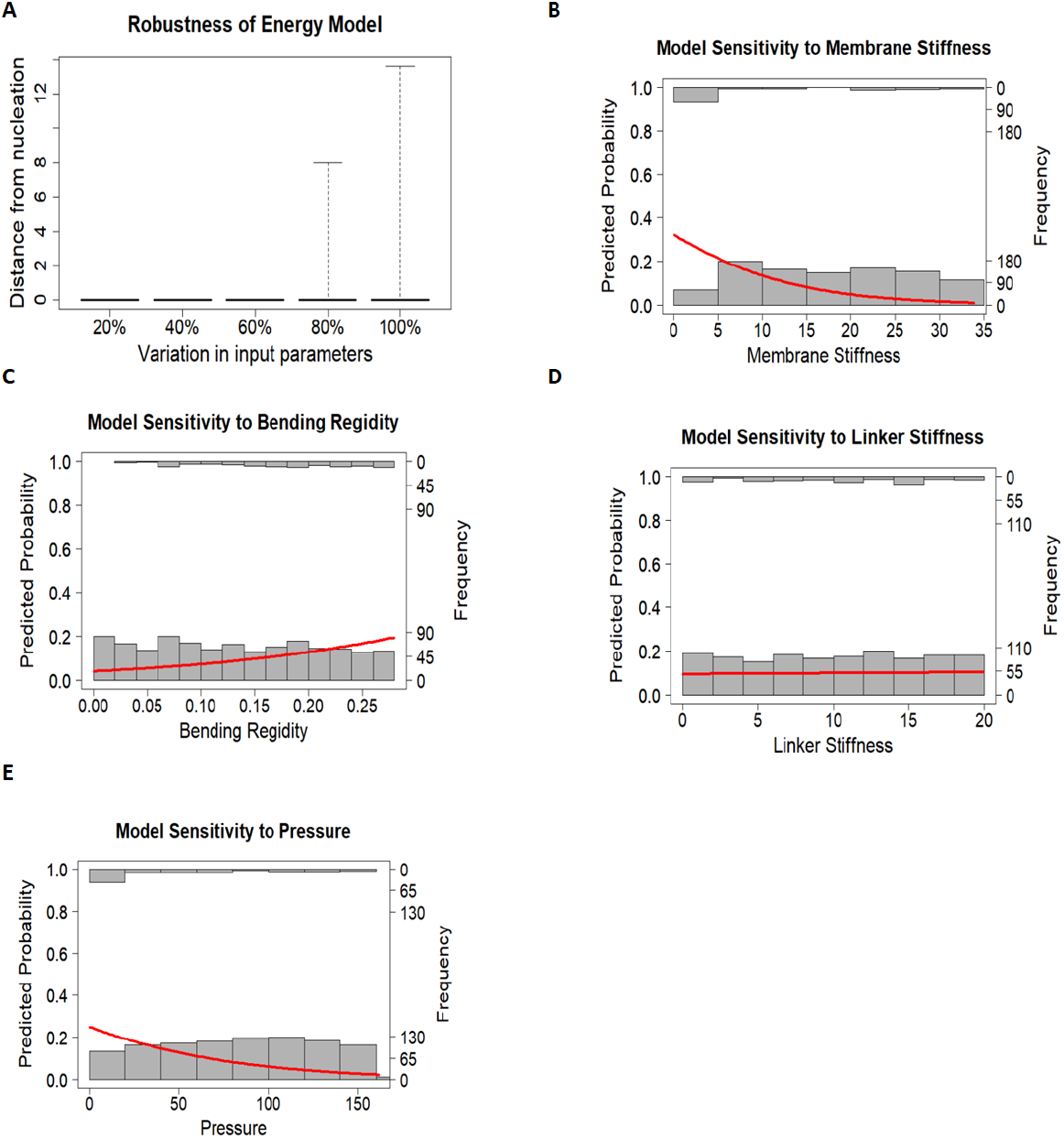
Robustness and sensitivity of the energy functional. 1000 uniform random vectors of the model parameters, *α, β,κ*, Π, are simulated and used to investigate the robustness of the energy functional. In A: a box plot shows the distribution of model output when model input parameters are varied by 40%, 60%, 80% and 100% of their measured values. B-E: shows histograms of model output across individual parameters at 100% variation as well as predicted probabilities from a logistic regression model (red curve). B: Model sensitivity increases with decreasing membrane stiffness coefficient, pvalue = 4.237280*e* – 15. C: Model sensitivity increases with increasing bending rigidity, pvalue = 1.539986*e* – 07. D: Model sensitivity is unaffected by linker stiffness, pvalue = 9.967809*e* – 01. E: Model sensitivity increases with decreasing ambient hydrostatic pressure, pvalue = 4.240195*e* – 10.

In over 70% of the blebs we examined, a distinct narrow gap was observed in the cortex scar - the part of the cortex from which the membrane first detached. An example is shown in Fig 4E1 as feature *d*. The gap at *d* is about at the midpoint between shoulder points *a* and *b* and opposite the furthest extent *c*. This gap is already present in the pre-bleb image as seen in Fig 4D1 and 4D2. The presence of the visible gap in the pre-bleb image distinguishes this gap from normal post bleb cortex disassembly. It associates the gap with bleb initiation. The gap at *d* lines up well with the nucleation point determined by furthest extent. Thus, we frequently used the presence of a gap in the early cortex scar to confirm the nucleation sites determined using furthest extent.

### Membrane tension and local hydrostatic pressure are critical forces driving bleb nucleation

In order to evaluate the contribution of each component of the energy functional to its accuracy, we eliminated each factor and asked how well the model predicts the nucleation sites. A total of 23 cells and 86 blebs were examined. Each row of Table 2 represents the result of eliminating one of the components of the energy functional. First, we eliminated membrane tension and lost all predictability (0.0%), suggesting that membrane tension is key to the efficacy of the model (Fig. 5B). Almost all model predictions were maintained when curvature (95.3%) and linker tension (98.8%) were each removed from the model. This suggests a minor role for these forces in determine where blebs nucleate (Fig. 5C,D). When the pressure component was removed, predictability was only 3.5%, suggesting huge importance (Fig. 5E). These results are striking. Membrane tension and local pressure are highly significant while curvature and linker protein tension appear to be largely irrelevant. It would appear that we could remove the latter two terms without vastly changing the effictiveness of the functional. One potential reason curvature appears less significant here is that we are focused on the bleb shoulder, a small region relative to cell boundary, where variation in curvature is small. Again the narrow region of interest explains why the linker protein tension appears insignificant.

It is especially important that membrane tension and variable pressure combine to yield an effective tool for locating bleb nucleation. Central to our energy model is the hypothesis that local contraction by myosin II in the blebbing region leads to local variation in hydrostatic pressure. Also, a major contributor to membrane tension is cortical tension which is regulated by myosin II and helps to prevent cytoskeletal collapse when the cell is subjected to high external load. Indeed, the level of myosin II in the cortex has been shown to increase in response to increasing levels of confinement [34]. Together, these results suggest a critical role for myosin II in bleb nucleation, supporting hypotheses made in previous studies [2, 8, 26, 34].

**Table 2.**
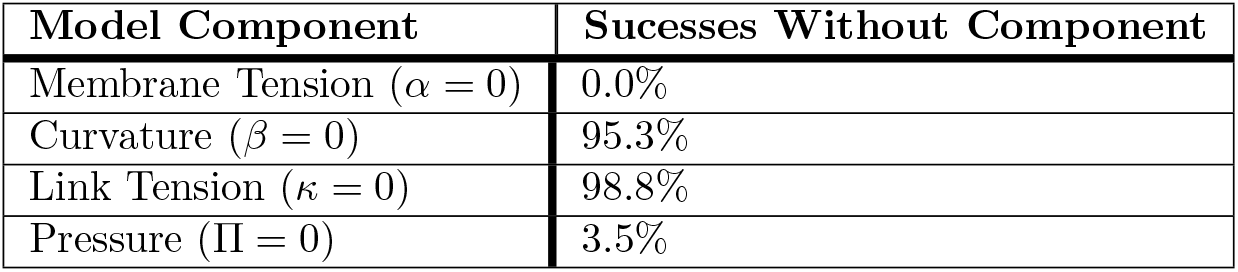
Significance of forces within the energy functional.

| Model Component | Sucesses Without Component |
| --- | --- |
| Membrane Tension ($\alpha = 0$) | 0.0% |
| Curvature ($\beta = 0$) | 95.3% |
| Link Tension ($\kappa = 0$) | 98.8% |
| Pressure ($\Pi = 0$) | 3.5% |

### Myosin II clusters form in locations of bleb formation

Our result on the role of myosin II, based on our mathematical model, is not biological in the usual sense as it discusses factors that are not observable. However, it identifies a protein of importance, myosin II, that can be studied using biological assays. In short, the mathematics has provided insight into the cell that is not routinely available. To verify our conclusions from the mathematical model, we examined the localization of myosin II in the cell cortex for *D. discoideum* cells actively blebbing while chemotaxing under an agarose gel. Fig 6A shows the microscopy image of a cell expressing myosin-GFP. We identify 6 non-posterior sites enriched with myosin II. These sites are also identified in the normalised florescence intensity profile in Fig 6C. We refer to these as myosin II clusters. Interestingly, after 4.28 seconds, a nascent bleb emerges from site 5 as shown in Fig 6B. We observe myosin II beginning to enter the nascent bleb, possibly to help retract the newly formed cortex. The emergence of blebs from myosin II cluters occured in about 80% of the cases we examined. This is remarkable as it provides biological evidence in support of an association between anterior clustering of myosin II and bleb initiation.

**Fig 6.**
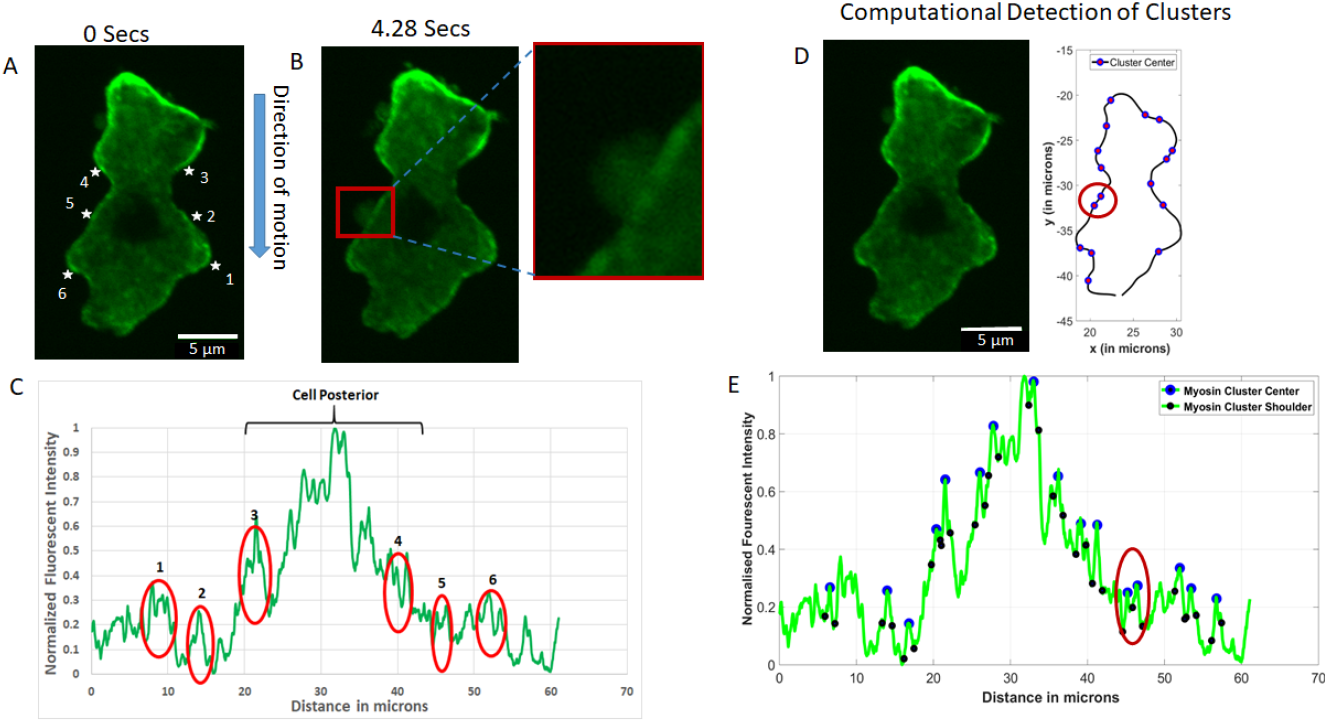
Microscopy evidence for the anterior clustering of myosin II prior to bleb initiation. The image shows myosin II tagged with a green fluorescent protein (GFP). A: Myosin II clusters, denoted 1-6, appear away from the cell posterior. B: A bleb emerges from cluster 5, 4.28 seconds later. Now the myosin II has invaded the newly formed bleb. C: Myosin II clusters are identified on the fluorescent intensity profile. D: Distribution of clusters around the cell boundary as identified by the cluster detection algorithm. E: Clusters identified on the intensity profile by the cluster algorithm.

To ensure that our observation of clusters was unbiased, we developed an algorithm (see materials and methods for details) that detects clusters directly from fluorescence intensity data. The results from this computational approach matched our observationally detected clusters (see Fig 6(D and E)).

## Discussion

While our observations and modeling correlate myosin clustering, cortex gaps and bleb initiation, no mechanistic explanation is readily apparent. However, examination of the literature suggests that gaps may be caused by myosin clustering [2,34,38]. Myosin has been observed to localize to regions of the cortex under greatest stress [34]. In the the presence of large numbers of myosin II, the mean persistence length of actin filaments has been found to reduce [39]. Shorter actin filaments are more likely to escape the cortical network [40], thereby weakening the overall structure. As stress on the cortex continues to ramp up from myosin II accumulation, the weakened cortex will exceed its elastic modulus and collapse [41,42].

Based on the evidence presented so far in this study and the stated results from literature, we propose the following model for bleb initiation that explains the role of cortex gaps. For cells chemotaxing in compressive 3D environments, the overall increase in membrane tension draws more myosin II from the cytoplasm to the cortex. In addition, the presence of cAMP gradient initiates the movement of myosin II to the cell anterior where it clusters at locations of high membrane tension (Fig 7A). The clustering has two effects. First, it reduces the mean persistence length of actin filaments thereby allowing shorter filaments to escape, leaving behind a weaker structure. Secondly, it increases strain on the weakened network through increased myosin II contractility. Beyond some critical strain, the network ruptures and detaches neighboring cortex/membrane linker proteins. Pressurized cytosol then flows through the created gap, detaches the membrane further and forms the bleb.

**Fig 7.**
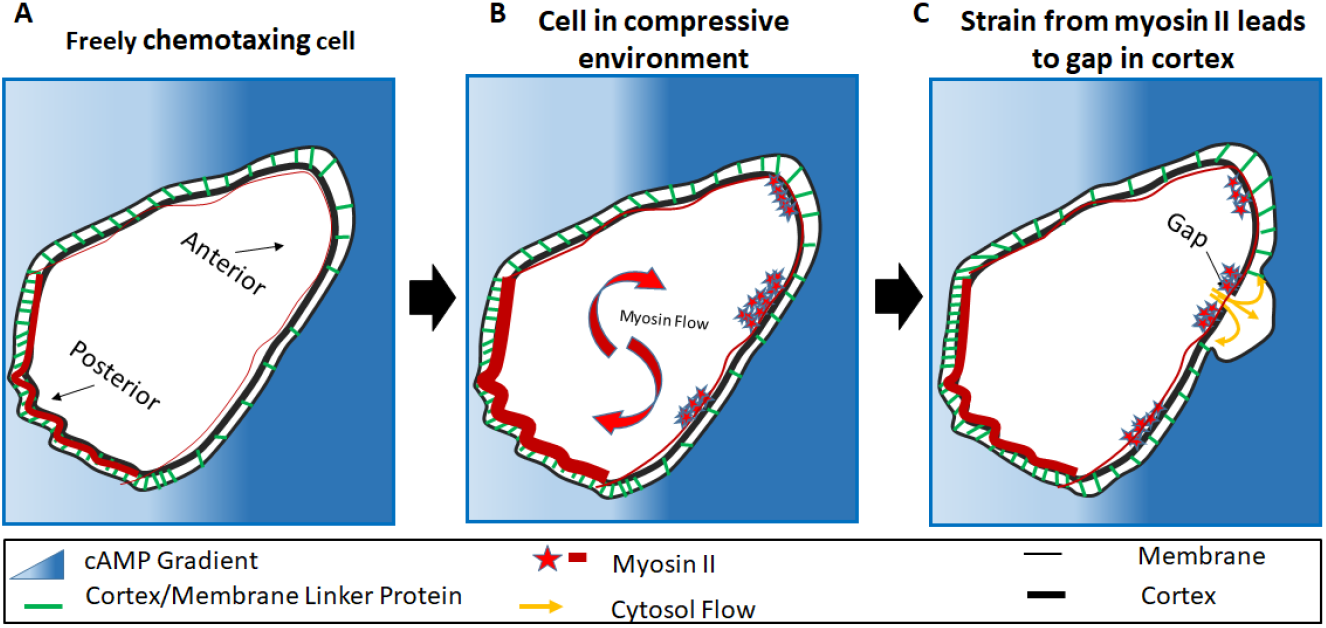
Model for bleb initiation. The image illustrates the cortex/membrane complex with connector protein distribution, posterior and anterior identified. A: The cell detects the cAMP gradient. B: Myosin II density increases due to compression from the environment. Myosin II is attracted by the cAMP causing a forward recruitment. it then forms clusters at the anterior. C: Myosin II contractions results in cortex degradation, a gap forms weakening the connection to the membrane. The bleb follows.

This model brings together two processes for bleb initiation that have previously been suggested (pressure differentials and cortex degradation). However, this is the first instance where mathematical modeling and experimentation have combined these two processes into one unified model of bleb initiation. This emphasized the critical role mathematics will continue to play in biological research.

## Acknowledgments

The authors thank Chandra Mangroo and Michael Barile for their assistance in data collection. We thank L. Leslie Liu for statistical support. This work was supported by grants to Derrick Brazill from the National Science Foundation (MCB-1244162), a PSC-CUNY grant (692710047), as well as Research Centers in Minority Institutions Program grants from the National Institute on Minority Health and Health Disparities (8 G12 MD007599) from the National Institutes of Health. Zully Santiago is supported by the Research Initiative for Scientific Enhancement (RISE) at Hunter College funded by NIH grant GM060665.

**Supporting information**

